# SIMMAM 3.0 – Updating the Toolbox for the Conservation of Marine Mammals

**DOI:** 10.1101/2022.03.14.484333

**Authors:** Alencar Cabral, André S. Barreto

## Abstract

SIMMAM (Sistema de Apoio ao Monitoramento de Mamíferos Marinhos) has been available for marine mammal researchers and environmental agencies in Brazil since 2005. However, it had to be updated, to adapt to new standards and data sharing policies. SIMMAM 3.0 uses more modern technologies (PHP 7.4, Symfony 5.x, PostgreSQL 11.x, PostGIS 3.x, Leaflet 1.7) and is compliant to DarwinCore (DwC) standard and able to exchange data with other data portals that use it. Its database now holds more than 44000 marine mammal records and is an important tool in the decision-making process of Brazilian environmental agencies.

## Resumo

O SIMMAM (Sistema de Apoio ao Monitoramento de Mamíferos Marinhos) está disponível para pesquisadores de mamíferos marinhos e agências ambientais no Brasil desde 2005. No entanto, sua atualização foi necessária para se adequar a novos padrões e políticas de compartilhamento de dados. O SIMMAM 3.0 usa tecnologias mais modernas (PHP 7.4, Symfony 5.x, PostgreSQL 11.x, PostGIS 3.x, Leaflet 1.7) e é compatível com o padrão DarwinCore (DwC) e capaz de trocar dados com outros portais de dados que também o utilizam. Seu banco de dados já possui mais de 44.000 registros de mamíferos marinhos e é uma importante ferramenta no processo de tomada de decisão dos órgãos ambientais brasileiros.

## 1. Introduction

Several worldwide initiatives, such as the Global Biodiversity Information Facility - GBIF have shown the importance of digital databases for the study and conservation of species. Marine mammals are a good example where these large-scale databases can contribute, as they naturally occur in low densities and usually in areas that are difficult to access. One of the main sources of information for marine mammals are stranded animals, as biological samples and can be obtained from them and thus they contribute not only with occurrence data but also with biological parameters. However, strandings are rare events and to be biologically meaningful they need to be accumulated over large distances, long times, or both.

To organize a database of marine mammal sightings and strandings along the Brazilian coast, the SIMMAM (Sistema de Apoio ao Monitoramento de Mamíferos Marinhos) project was started. It began as an internal project with collaboration between the *Laboratório de Oceanografia Biológica* (Biological Oceanography Laboratory) and the *Laboratório de Computação Aplicada* (Applied Computing Laboratory, presently *Laboratório de Informática da Biodiversidade e Geomática – LIBGeo*), that began using historical data, as well as sighting data collected by scientific observers from research projects coordinated by UNIVALI researchers. Afterward, an agreement for technical cooperation was signed with the Centro Mamíferos Aquáticos, at that time a part of IBAMA and now of ICMBio Its initial implementation has already been described [Moraes, 2005; Barreto et al., 2006], and was based on PHP 5.0 and Adobe Flex 3.0. Considering the development of new technologies and communication protocols, SIMMAM had to evolve to adapt to this new environment. This paper describes its new implementation and the evolution of its database in terms of contributions from different parties.

## 2. Technologies

The speed with which the technologies have changed in the last decades is ever increasing, not only in computational tools but also regarding communication protocols, data sharing policies, and data availability. To attend the needs of researchers and Brazilian environmental agencies that use the system, SIMMAM was almost completely rewritten using modern technologies, both internally as well as for the organization and implementation of a communication protocol that made SIMMAM compatible with existing data aggregators.

SIMMAM 3.0 now conforms to the DarwinCore (DwC) standard, which is an international scientific initiative of the Taxonomic Database Working Group - TDWG. The data architecture adopted is compatible with GBIF, which will allow SIMMAM to become a data publisher of marine mammal occurrences.

For the development of SIMMAM 3.0 it was decided to maintain the premise of using free source code tools. The development of the system followed the MVC methodology. On the server side, PHP 7.4 was used with Symfony 5.x framework. For the web client side, we opted to render the site on the server side and deliver an HTML + JavaScript page with Bootstrap 5 to the browser (Figure 1). The data exchange API was also implemented in PHP, following the XML standard of DwC.

**Figure 1.**
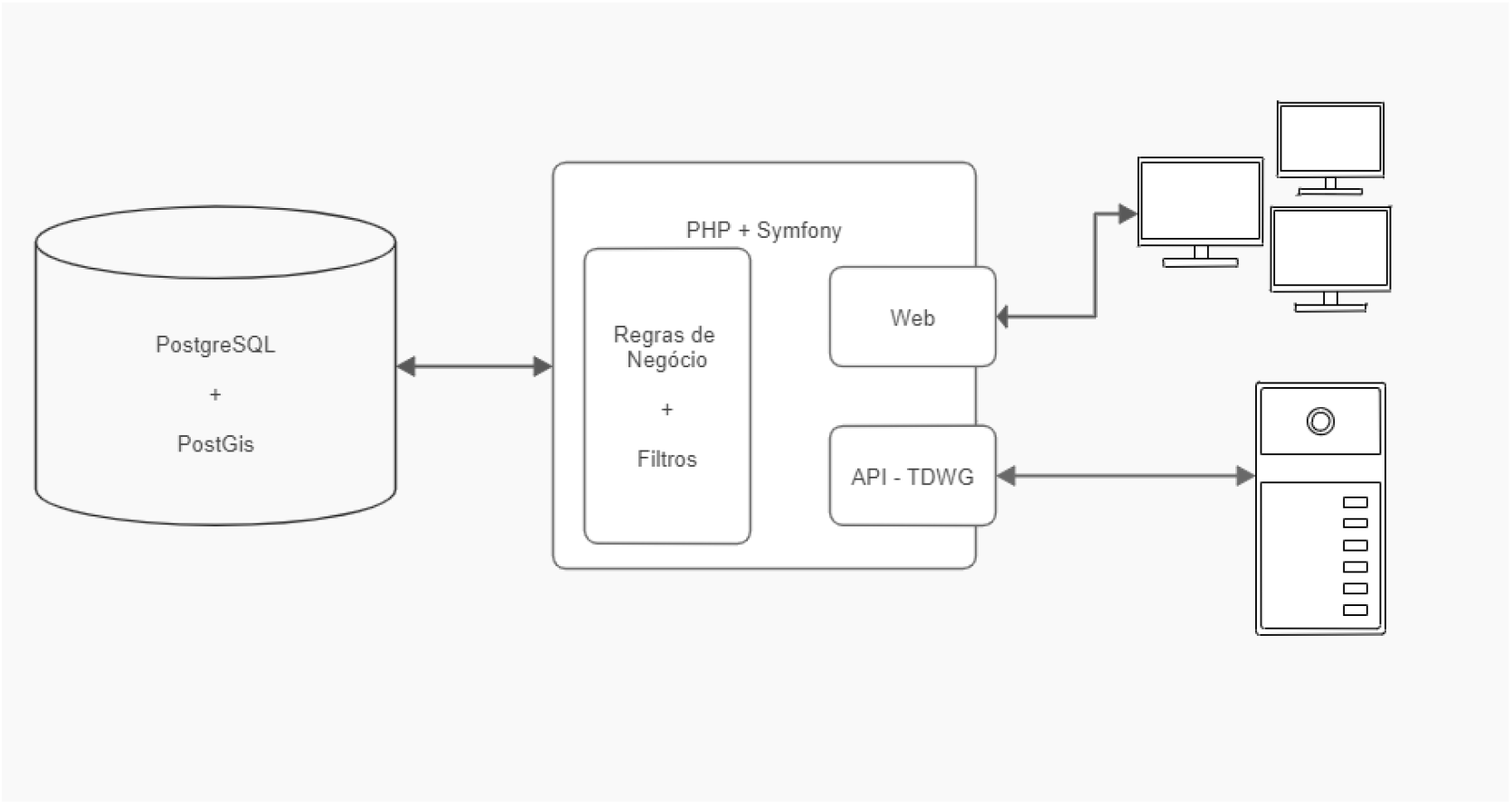
Simplified block diagram for SIMMAM 3.0.

Data is stored in PostgreSQL 11.x with PostGIS 3.x which allows manipulation of geospatialized data. The tables were structured according to the DwC standard (Figure 2), to reduce the complexity of the communication API.

**Figure 2.**
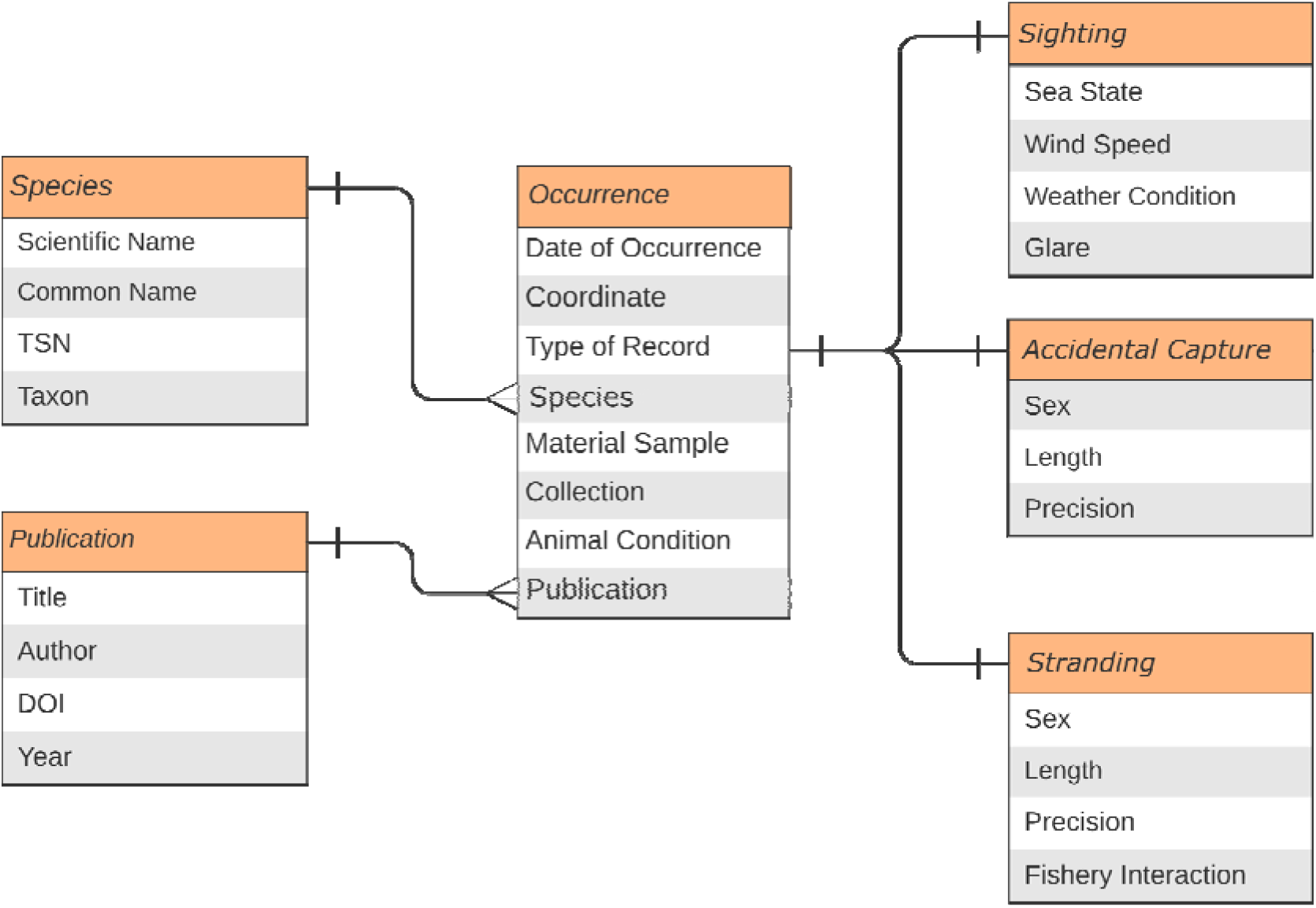
Relationship diagram for the main entities in SIMMAM 3.0.

To ensure the security of private data, a business rule was implemented in such a way that all data is compartmentalized, preventing private data from being seen by other users. The only exception is for users in the ‘Environmental Agency’ category who can see all the data, as they need to have access to all occurrences for decision making purposes.

As all occurrences in SIMMAM need to have a geographic position, the main interface for users to view the data is through an interactive WebGIS. The WebGIS implemented has filters by taxon and by type of occurrence (sighting, stranding, incidental capture) to allow users to focus on specific groups of organisms or kinds of data. To better display areas with high density of records, the occurrence layer was clustered, grouping and ungrouping records according to the zoom level (Figure 3). This allows the user to view hotspots of occurrences without generating visual clutter. In the webGIS construction, Leaflet Map [Agafonkin, 2020] was used with the OpenStreetMap base map, as it is a modern map engine, has functionalities optimized for mobile devices, and does not have any external dependency. Leaflet supports multiple layers and is compatible with the Open Geospational Consortium (OGC) standard such as support for map mosaics, georeferenced images, WMS [Leaflet, 2020] and GeoJSON [IETF, 2021].

**Figure 3.**
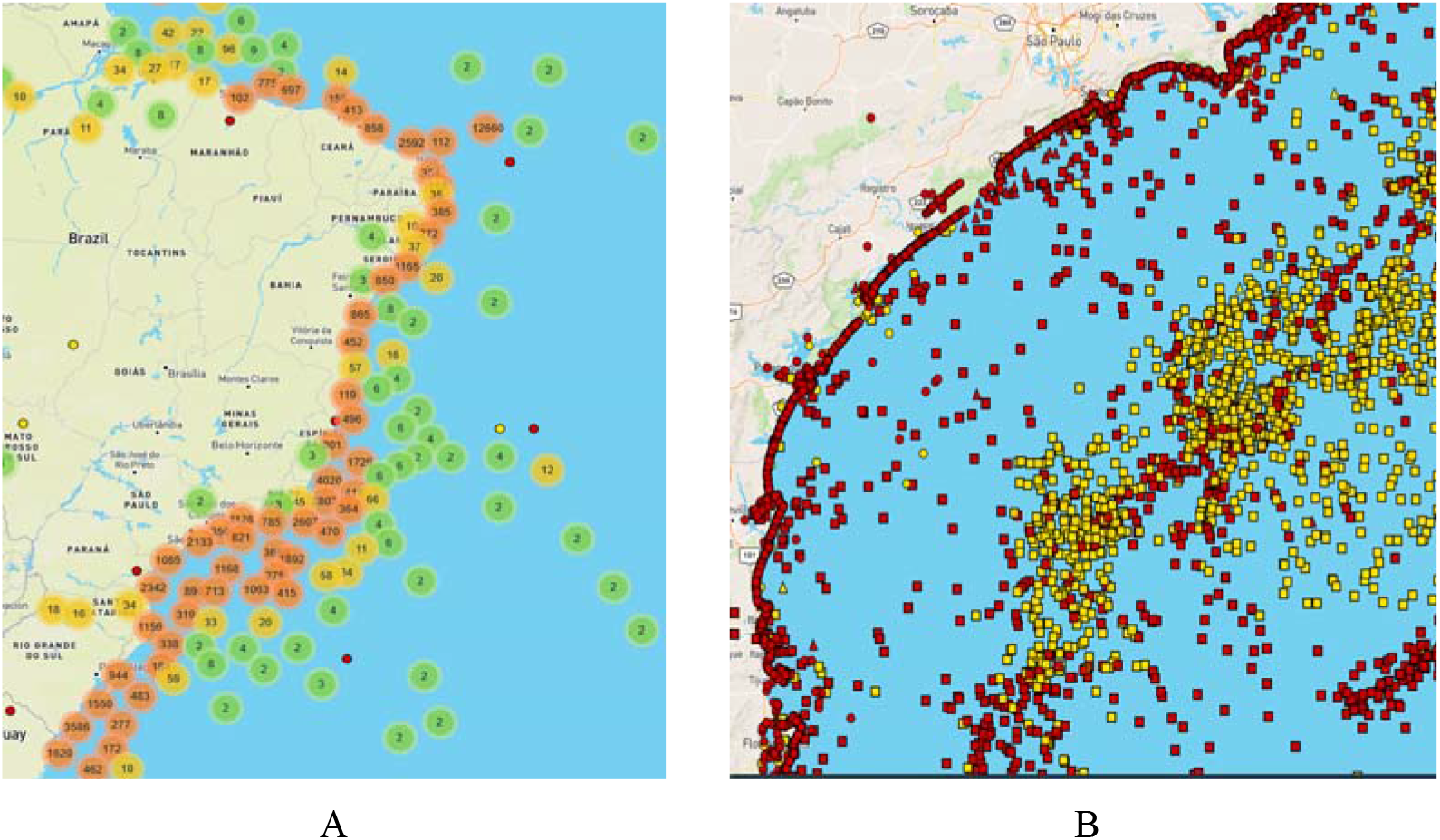
Example of the WebGIS implemented in SIMMAM 3.0, showing data (A) aggregated in groups at low zoom levels, and (B) ungrouped with higher zoom levels. Yellow points represent public data, and red points private data.

However, a WebGIS alone would not be enough to manipulate SIMMAM data, thus it is possible to export data in spreadsheets (xls format). The exported spreadsheet has all the information that the user inserted for each type of record, such as species, geographic position, date, group size, etc.

One key aspect of biological information is the taxonomic identification. Due to advances in the field, organisms previously identified with a name (family, genus, species) can have this name changed. To avoid taxonomic instability in SIMMAM, it uses the taxonomic list provided by the Integrated Taxonomic Information System – ITIS (www.itis.gov). It aims to provide reliable information on species names and their hierarchical classification, with curatorship of the species listings to keep them up to date. As the taxonomic classification of mammals is very stable, it was decided to keep a copy of the ITIS database locally to reduce latency, being updated on demand.

The types of occurrence records currently supported by SIMMAM are stranding, accidental capture and sightings. All these occurrence records have fields for defining the best taxonomic level, geo-referencing the occurrence, information on biological material collected and the responsible for the data. Stranding and accidental capture records contain information regarding the state of the animal (alive or dead), the condition of the carcass (decomposition stage), sex and length. For sightings it is possible to inform environmental parameters such as weather condition, sea state, wind speed, as well as if it was a single animal, part of a group and group size.

## 3. Implementation and Use by the Scientific Community

The first version of SIMMAM was made available to the Centro Mamíferos Aquáticos - CMA, and in 2007 it started to use SIMMAM as the main tool to integrate data of the different institutions that composed the Brazilian Stranding Network of Aquatic Mammals (*Rede de Encalhes de Mamíferos Aquáticos do Brasil*, REMAB). On the same year, SIMMAM was presented to the then General Coordination of Oil and Gas (*Coordenação Geral de Petróleo e Gás* – CGPEG), current General Coordination of Marine and Coastal Enterprises (*Coordenação-Geral de Licenciamento Ambiental de Empreendimentos Marinhos e Costeiros* – CGMAC), which identified the possibility of using it to aggregate and organize marine mammal sighting data generated by marine mammal observers [Barreto et al., 2019]. A cooperation agreement was signed between UNIVALI and IBAMA, for the review of reports between 2000 and 2008, organizing and inserting all marine mammal data in SIMMAM (BRITTO; 2009). After that, sighting data was uploaded to SIMMAM directly by the licensed companies. In 2010 a research grant was awarded to SIMMAM as part of the SISBIOTA initiative of the Brazilian government. This allowed to strengthen the network of scientific institutions that contributed data to SIMMAM, resulting in an increase in the inclusion of data in 2013, when this grant finished (Figure 4).

**Figure 4.**
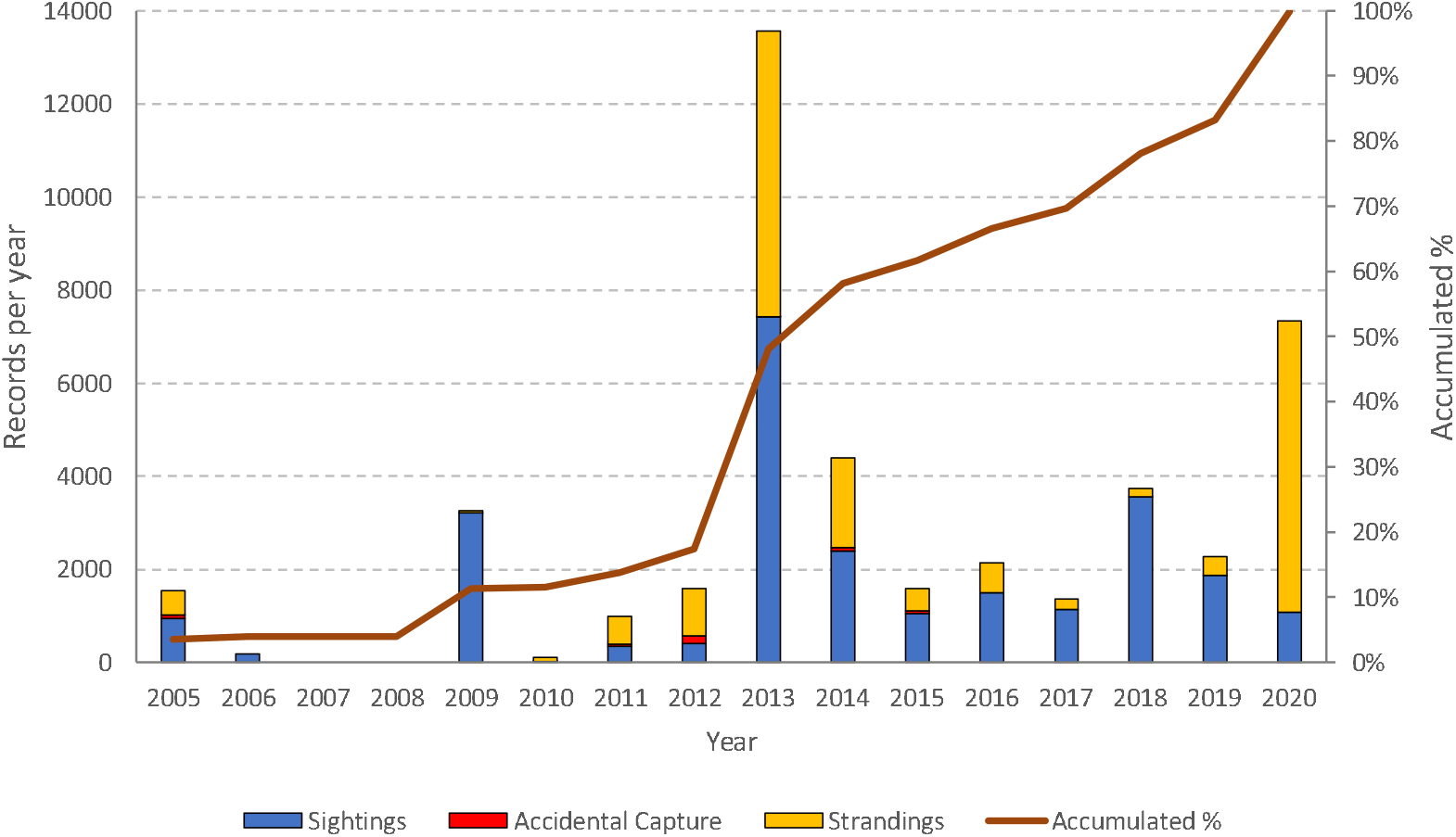
Temporal evolution of data included in SIMMAM, considering the different types of records.

As of December 2020, SIMMAM had 125 active users and held 44,096 aquatic mammal records. Of these, 61% records are private, but this proportion is very different depending on the type of record (Figure 5). For strandings, that are in most part submitted by research institutions, 91% are private as they are the results of individual efforts. But for sightings 61% are public, as they come mostly from the oil industry as part of the environmental licensing of their operations, and they mirror the public reports that have been delivered to IBAMA. As mentioned before, all the data held in the SIMMAM database, regardless of its public availability, can be seen by Brazilian environmental agencies (IBAMA and ICMBio).

**Figure 5.**
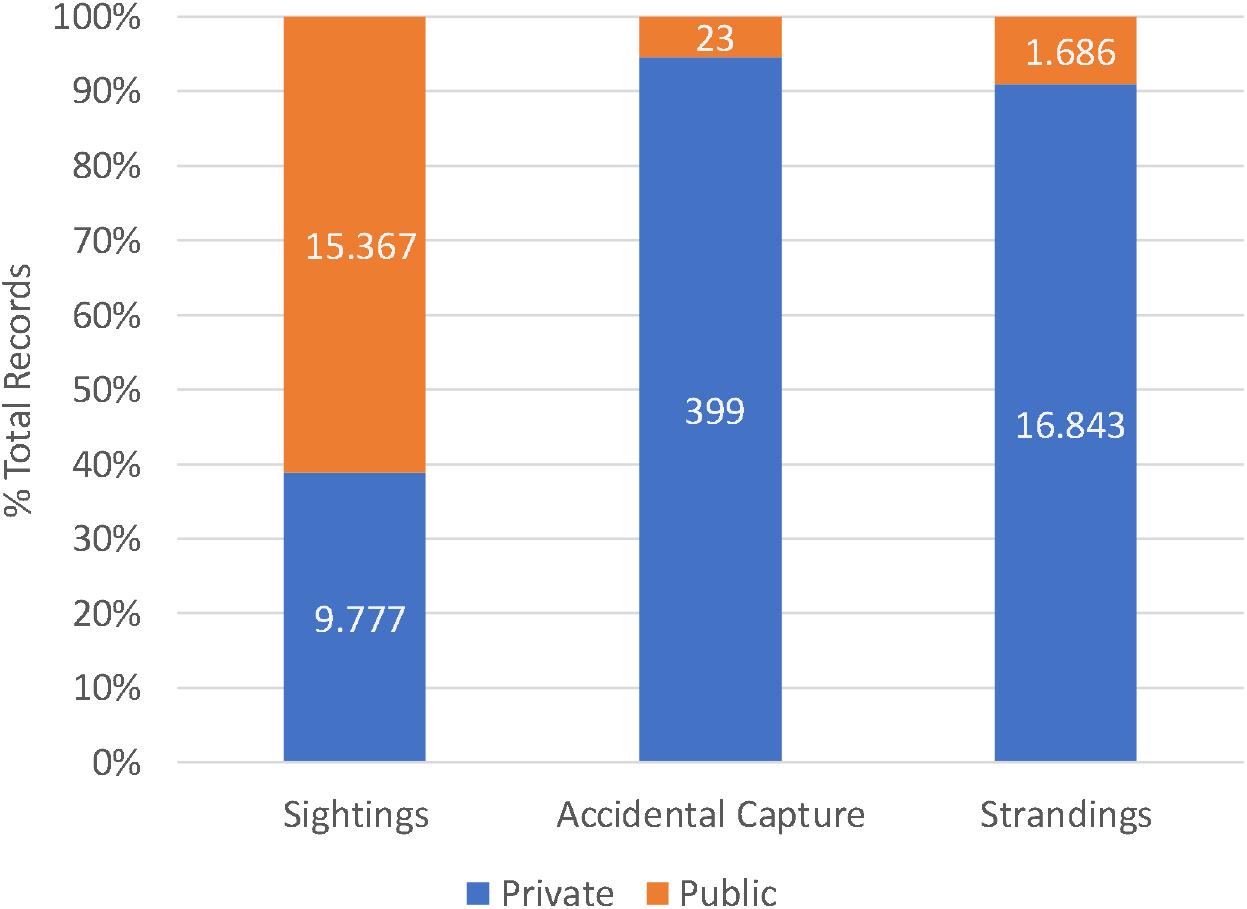
Data in SIMMAM 3.0, separated by type of record (sightings, strandings, accidental capture) and availability (private, public).

The option to allow government agencies to use the whole dataset is extremely important for management purposes, as it enables the environmental agencies to use even unpublished data generated by research institutions. But as the data is not available for the general public, it does not compromise their future use in academic publications. Also, the limited visualization of private data in the WebGIS (red dots in Figure 3), where details of the record such as species and date are not shown, serves as an indication for other researchers that a specific institution has data on marine mammals in a specific area, fostering collaborations among institutions.

## References

Agafonkin, V. (2020) “Leaflet”. https://leafletjs.com, 2020

Barreto, A. S.; Cabral, A.; Taufer, R. M.; Almeida, T. C. M.; Owens, A. L. (2019) “Conhecimento sobre mamíferos marinhos gerado pela indústria de sísmica através do sistema de apoio ao monitoramento de mamíferos marinhos (SIMMAM)”, In: Barbosa, A. F.; Owens, A. L.. (Org.). IBAMA e Indústria de Pequisa Sísmica: em busca de conhecimento e sustentabilidade através do licenciamento ambiental. 1ed. Rio de Janeiro: Mind Duet Comunicação e Marketing, p. 102–114.

Barreto, A. S.; Moraes, C. G.; Sperb, R. M. and Bugghi, C. H. (2006) “Using GIS To Manage Cetacean Strandings”. Journal of Coastal Research, SI 39, 1643–1645.

Britto, M. K. (2009) “Mamíferos Marinhos, a Atividade de Prospecção Sísmica e o Uso do Sistema de Monitoramento de Mamíferos Marinhos – SIMMAM”, 118 p. Master’s Thesis (Mestrado em Mestrado em Ciência e Tecnologia Ambiental) - Universidade do Vale do Itajaí.

IETF (2021) “The GeoJSON Format – RFC 7946”https://datatracker.ietf.org/doc/rfc7946/

Leaflet.wms (2020) “An all-in-one WMS plugin for Leaflet”. https://github.com/heigeo/leaflet.wms

Moraes, C. G. (2005) “SISTEMA DE MONITORAMENTO DE MAMÍFEROS MARINHOS - SIMMAM: uma ferramenta para o estudo de avistagens e encalhes na costa brasileira”. 69 p. Master’s Thesis (Mestrado em Mestrado em Ciência e Tecnologia Ambiental) - Universidade do Vale do Itajaí.

